# Neural substrates underlying the expectation of rewards resulting from effortful exertion

**DOI:** 10.1101/2024.02.01.578411

**Authors:** Aram Kim, Vikram S. Chib

**Affiliations:** Department of Biomedical Engineering, Johns Hopkins School of Medicine; Baltimore, MD 21205, USA; Center for Movement, Kennedy Krieger Institute; Baltimore, MD 21205, USA; School of Exercise and Nutritional Sciences, San Diego State University, San Diego, CA, 92104, USA; Kavli Neuroscience Discovery Institute, Johns Hopkins University; Baltimore, MD 21205, USA

**Keywords:** Effort, Decision-making, Ventral striatum

## Abstract

Decisions to exert effort for reward are often conceptualized as reflecting a trade-off between anticipated benefits and effort costs. However, economic theories propose that valuation is reference-dependent, such that behavior is shaped relative to expected outcomes rather than absolute reward magnitude. Whether reward expectations influence effort expenditure through such reference-dependent mechanisms, and how these computations are represented in the brain, remains unclear. Using an incentivized grip-force task during functional magnetic resonance imaging, we experimentally manipulated reward expectations independently of reward outcomes. Behavioral results showed that effort exertion varied systematically with reward expectations, indicating that individuals evaluated exertion relative to a reference point. Neural activity in the ventral striatum tracked both prospective reward expectations and deviations between expected and realized earnings. Moreover, these neural signals predicted individual differences in reference-dependent modulation of effort. Together, these findings identify ventral striatal representations associated with reference-dependent effort valuation and suggest that reference-dependent computations contribute to motivated behavior.

**Significance statement:** Decisions to exert effort are commonly described as reflecting a trade-off between reward and effort cost. However, whether effort valuation is shaped by reward expectations rather than absolute incentives remains unclear. Using functional magnetic resonance imaging and an incentivized grip-force task, we show that reward expectations serve as a reference point that systematically influences effort expenditure. Neural signals in the ventral striatum tracked both reward expectations and deviations between expected and realized outcomes, and these signals predicted individual differences in reference-dependent effort behavior. These findings suggest that reference-dependent valuation contributes to the neural control of motivated action.

## Introduction

Reward expectations influence decisions to engage in effortful behavior. Field studies have shown that willingness to exert effort is shaped not only by the incentives offered but also by expectations about prospective earnings and prior outcomes (Camerer et al., 1997; Fehr and Goette, 2007; Farber, 2008; Crawford and Meng, 2011; Gill and Prowse, 2012; Allen et al., 2014; Bartling et al., 2015; Gutierrez et al., 2021).

For example, taxi drivers adjust their working hours according to their expected daily income (Camerer et al., 1997; Crawford and Meng, 2011), and professional athletes increase their effort when prior experience leads them to expect success (Bartling et al., 2015). These findings are consistent with theories of reference-dependent valuation, which propose that behavior is evaluated relative to an internal reference point rather than absolute outcomes alone (Kahneman and Tversky, 1979; Kőszegi and Rabin, 2006; Fehr and Goette, 2007; Gneezy et al., 2017; O’Donoghue and Sprenger, 2018). Although reference dependence has been extensively studied in economic choice, comparatively little is known about how reward expectations influence neural computations underlying effortful exertion.

Neurobiological studies of effort-based decision-making have primarily focused on how the brain integrates anticipated rewards with the costs of exertion to guide motivated behavior (Pessiglione et al., 2007; Croxson et al., 2009; Prévost et al., 2010; Kurniawan et al., 2011, 2013; Chong et al., 2017; Hogan et al., 2019, 2020; Dryzer et al., 2025; Steward et al., 2025). This work has implicated corticostriatal circuits in representing effort costs, reward value, and willingness to exert effort. However, most studies have examined effort valuation in terms of absolute incentive magnitude, despite evidence that valuation in other domains is strongly shaped by contextual expectations and reference points. Whether reward expectations similarly alter neural computations underlying effort exertion remains unclear.

Previous neuroimaging studies have implicated the ventral striatum (VS) in multiple computations relevant to reference-dependent valuation. Activity in the VS has been associated with subjective value representations (Kable and Glimcher, 2007; Tom et al., 2007; Chib et al., 2012, 2014; Levy and Glimcher, 2012; Bartra et al., 2013; Dunne et al., 2019) and reward prediction errors reflecting deviations between expected and realized outcomes (O’Doherty et al., 2003; Pessiglione et al., 2006; Rutledge et al., 2010; Daw et al., 2011). In addition, VS activity has been linked to reference-dependent valuation during economic exchange decisions (Martino et al., 2009). Together, these findings suggest that the VS may contribute to computations through which reward expectations shape effort expenditure. However, whether VS activity encodes reference-dependent signals that influence effort exertion has not been directly tested.

Here, we investigated the neural mechanisms through which reward expectations influence effort exertion. Using an incentivized grip-force task during functional magnetic resonance imaging, we experimentally manipulated effort-based reward expectations independently of reward outcomes. We hypothesized that reward expectations would serve as a reference point that modulates effort exertion, such that greater expected rewards would increase willingness to exert effort. We further hypothesized that VS activity would track deviations between expected and realized earnings, consistent with a role for the VS in effort-based reference-dependent valuation.

## Materials and Methods

### Participants

34 healthy, right-handed young adults (17 female, mean age 24 ± 7 years) participated in the study. Four participants were excluded from the behavioral analysis due to 1) a lack of understanding of effort-reward associations during either the *Training* or *Recall* phases (correlation coefficient < 0.7) and 2) more than 50% failed trials during the *Effort Provision* phase. Three additional participants were excluded from the neuroimaging analysis due to excessive head movement. Excessive head movement was defined as 1) mean framewise displacement (FD) greater than 1 mm or 2) more than 43 outlier frames (determined by FD greater than 1 mm), corresponding to approximately 2 minutes of the 11-minute scan. Therefore, the final behavioral and neuroimaging analyses included 30 and 27 participants, respectively. We prescreened all participants to exclude those with a prior history of neurological or psychiatric illness. The Johns Hopkins School of Medicine Institutional Review Board approved this study, and all participants provided written informed consent.

### Experimental Setup

Presentation of visual stimuli and data acquisition for the experiment were achieved using custom MATLAB 2020b (MathWorks, Natick, MA) scripts using the *Psychtoolbox* Version 3 (Kleiner et al 2007). A hand clench dynamometer (TSD121B-MRI, BIOPAC Systems, Inc., Goleta, CA) was used to record the exerted grip force. Force signals were amplified 1000 times (DA100C, BIOPAC Systems, Inc., Goleta, CA) and sampled at 1,000Hz (NI-DAQ card, Model PCIe-6321, National Instruments, Austin, TX).

### Experimental Design

Prior to the experiment, we informed the participants that they would receive a fixed show-up fee of $25 and earn additional money based on their performance on 10 randomly selected trials at the end of the fMRI session. Total earnings were revealed to participants at the end of the experiment to minimize the effect of historical earnings on subsequent performance during the experiment.

#### Reference-Dependent Exertion Task

When participants arrived at the laboratory, they read an experiment instruction package that guided them through the fMRI session. The fMRI experiment began with the acquisition of participants’ maximum voluntary contraction (MVC) (Fig. 1A). They held the hand-clench dynamometer in their dominant hand and, when cued, squeezed as hard as possible for three consecutive repetitions. Their MVC was calculated as the average of the three exertions, and the force level was converted into effort, defined as force relative to MVC, with 0 corresponding to no exertion and 100 to 80% of the participant’s MVC.

**Fig. 1.**
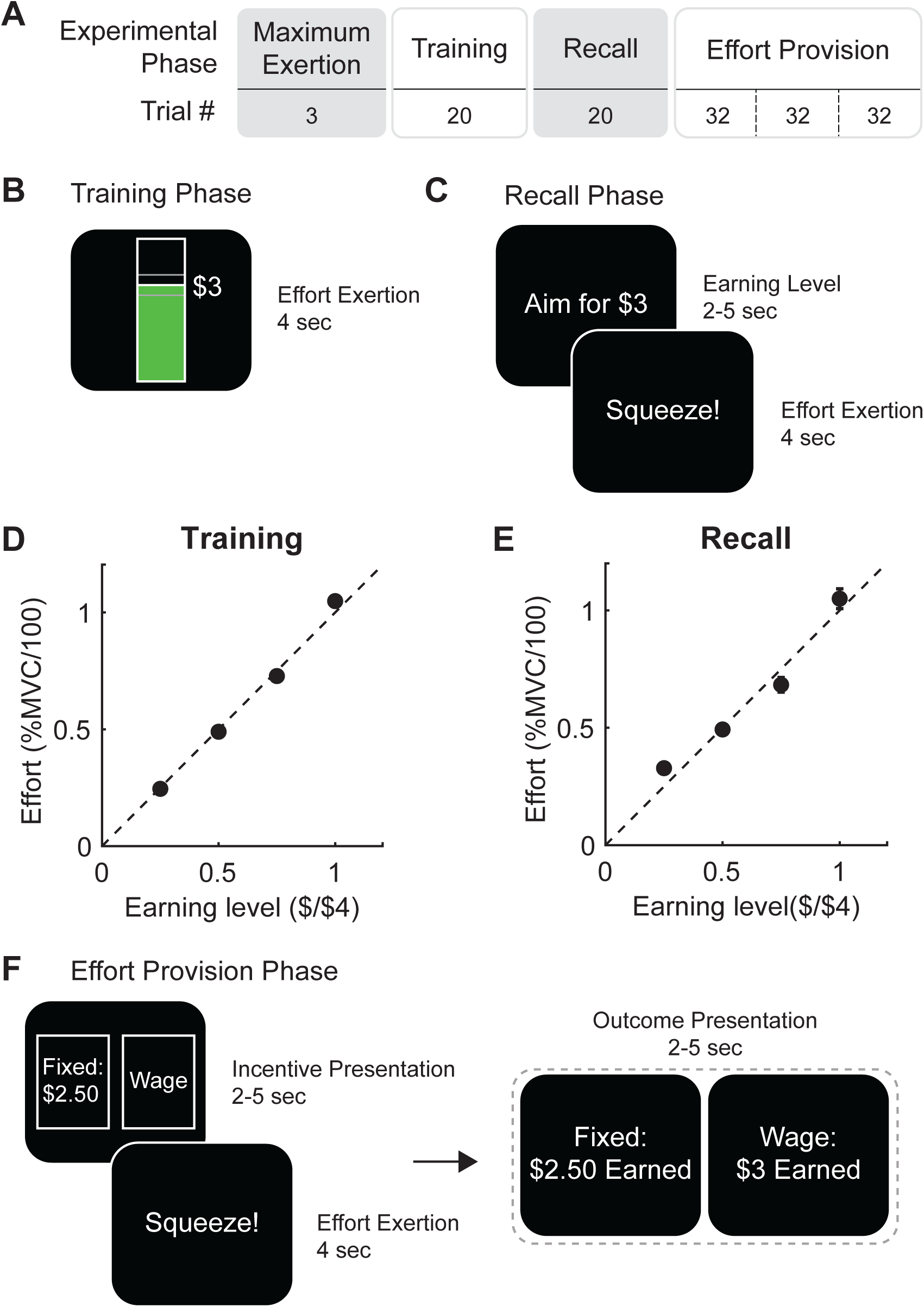
Experimental design. (**A**) Experimental schedule. Participants began the experiment by performing a maximum grip exertion, which was measured to normalize each individual’s strength for the effort task. They then performed a Training Phase in which they developed associations between the exertion required to obtain different earnings levels. (**B**) A training trial presented a target earning level, and a participant performed an effortful grip with real-time visual feedback of the exerted force represented as a bar that increased in height with increased exertion. The bar turned green when the participant reached and held the target exertion level. Effort levels ranged from $0 (no effort) to $4 (80% of maximum grip exertion). A Recall Phase (**C**) was performed to determine whether participants developed salient associations between exertion levels and earnings. During trials, participants were presented with a target incentive level and instructed to exert effort at that level without feedback of exertion. (**D**) Participants were able to accurately exert the target earnings levels during the training phase. (**E**) During the Recall Phase, participants exerted the appropriate effort associated with the presented earning level. (**F**) During each trial of the Effort Provision Phase, participants were presented with a risky option that would yield either a fixed monetary payment, regardless of the effort they exerted, or a wage payment proportional to the effort they exerted. Each option had an equal probability of occurring, and the fixed payment ranged from $0 to $4 to create different expectations among participants. The actual outcome, either the wage or fixed payment, was realized after individuals had a chance to exert effort.

After acquiring participants’ MVC, they completed a Training phase to develop an understanding of the associations between their effortful exertion and wage payment, which was proportional to their exertion (Fig. 1B). This phase consisted of 20 trials, with 5 trials for each target payment from $1 to $4, in $1 increments, presented in a randomized order. Each trial began with a fixation cross (2-10 s) and a Get Ready screen (1-2 s), followed by an effort exertion trial (4 s) featuring an interactive vertical bar that filled with red as participants gripped the hand dynamometer to reach the target range (defined as ±2.5 effort levels of the target, corresponding to $0.25), at which point the bar turned green. Participants were instructed to reach the target range as quickly as possible and to maintain the bar within it until the bar disappeared and the trial ended.

Next, participants completed a *Recall* phase to assess their understanding of the mapping between effort levels and corresponding wage earnings (Fig. 1C). In this phase, participants viewed a target wage-earning level for 2-5 s before the grip cue and received no online feedback on their exertion. They were also not shown the outcome of a trial (i.e., whether they successfully achieved the targeted wage-earning level with their exertion). The number of trials was the same as in the *Training* phase.

Lastly, participants performed an *Effort Provision* phase consisting of 3 blocks of 32 trials each (Fig. 1D). During the Incentive Presentation, each trial presented two possible outcomes: 1) Fixed payment and 2) Wage payment, each with a 50/50 chance of occurring. The fixed payment varied from $0.25 to $4 in $0.25 increments and was not contingent on a participant’s exertion. The wage payment was a piece-rate proportional to the amount of effort exerted. Participants did not actively select a single payment option but rather gripped the dynamometer to the level of effort they were willing to exert, considering the potential fixed payment (*Effort Provision*). Following exertion, one of the two payments was selected at random, and the outcome was revealed to participants. Participants were instructed to grip the dynamometer as quickly as possible and as hard as they saw fit to achieve their desired piece-rate earnings, given the fixed option presented. We considered a trial as a missed trial if participants failed (1) to initiate gripping quickly enough (within 1 s) or (2) to hold their grip exertion within 2.5 standard deviations for the last three seconds. Missed trials were not included in our final analyses. If the wage option was selected at random, piece-rate earnings were calculated based on the last 2 seconds of effort during the exertion phase; if the fixed option was selected, the fixed amount was paid out. Participants were instructed that 10 trials were randomly selected for additional payout (mean payout: $26.35 ± 6.72).

#### MRI Protocol

Images were acquired with a 3 Tesla Philips Achieva Quasar X-series MRI scanner and radiofrequency coil at the F.M. Kirby Research Center for Functional Brain Imaging at the Kennedy Krieger Institute. We collected high-resolution structural images using a standard magnetization-prepared rapid gradient-echo (MPRAGE) pulse sequence, providing full brain coverage at 1 mm × 1 mm × 1 mm.

Functional images were acquired using an echo-planar imaging (FE EPI) pulse sequence (TR = 2800 ms, TE = 30 ms, FOV = 240 mm, flip angle = 70°) at a 30° angle relative to the anterior commissure–posterior commissure (AC–PC) axis, which reduced signal dropout in the orbitofrontal cortex. Forty-eight slices were acquired at 3 mm × 3 mm × 2 mm, providing whole-brain coverage. We obtained five runs of functional images, consisting of *Training*, *Recall*, and three *Effort Provision* phases.

### Data Analysis

#### Behavioral Reference-Dependent Exertion Analysis

We tested the hypothesis that individuals would exert effort (*E*) relative to a fixed payment (*F*) using a linear mixed-effects model (LME). We also included potential confounding effects of (1) fatigue, (2) history of payment outcome, R(t-1), (3) history of outcome type (piece-rate/fixed), O(t-1), and (4) history of effort exertion, E(t-1) using as independent variables the trial number, reward magnitude in the previous trial, outcome type in the previous trial (fixed or wage payment), and magnitude of effort exertion in the previous trial, respectively to confirm that effort was driven by the exogenous referent (fixed payment) rather than by other potential confounding effects (Supplementary Figure 1).

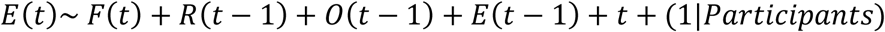

#### Image Processing

We used the SPM12 (Wellcome Trust Centre for Neuroimaging, Institute of Neurology, London, UK) software package to analyze the fMRI data. Functional images were 1) slice-timing corrected, 2) realigned to the first image to reduce head-motion effects, 3) normalized to the Montreal Neurological Institute (MNI) template, and 4) smoothed with a 3D Gaussian kernel (5 mm at full width at half maximum) to account for anatomical differences between participants.

The primary aim of the neuroimaging analysis was to identify brain regions that encode preferences for effort exertion relative to a reference point. Thus, we generated first-level, voxel-wise statistical parametric maps for each participant using general linear models (GLMs) with three conditions (*Incentive Presentation, Effort Exertion, and Outcome Presentation*). The *Incentive Presentation* epoch was modeled as a block, with a parametric modulator included for the expected value (0.5 × Fixed payment + 0.5 × Piece-rate payment) of a given trial. *Effort Exertion* was also modeled as a block and included a parametric modulator corresponding to a participant’s exertion level.

*Outcome Presentation* was modeled as an event and included a parametric modulator of the difference between a trial’s actual outcome and its expected value. We also included missed trials and outlier trials (mean effort ± 2 × standard deviation of effort) as separate nuisance regressors. Head motion parameters derived from the affine part of the realignment procedure during preprocessing were also modeled as separate nuisance regressors. In addition, we calculated FD based on the head motion parameters as an additional nuisance regressor (Power et al., 2014). If any frames had FD of 1 mm or higher, we created temporal masks of the data to specify which frames to ignore during analysis. These temporal masks were included as nuisance regressors. Finally, we created contrasts with the above parametric modulators for each GLM per participant.

#### Statistical Inference

We analyzed the VS signals within an independent region of interest (ROI) defined using Neurosynth.org (Yarkoni et al., 2011) with the search term “*value”* (peak MNI coordinates: left center = [-6 10 -4], right center = [12, 12, -8]). This ROI was comprised of bilateral VS. VS clusters from whole-brain contrasts shown in the Results were displayed at p < 0.005 in red with a 10-voxel extent threshold. We performed statistical inference in SPM using a small-volume correction (SVC) for multiple comparisons within the bilateral VS ROI described above at p < 0.05. To visualize signal patterns, we plotted effect sizes at the coordinates of peak activity. However, these plots were used solely for visualization and not for statistical inference.

#### fMRI Moderated Mediation Analysis

We conducted a multilevel moderated mediation analysis to examine whether trial-by-trial neural responses mediated the relationship between instantaneous fixed payment and effort exertion and whether this mediation was modulated by individual differences in the fidelity of individuals’ wage- effort association during the Recall Phase. Trial-wise neural estimates were obtained using a least-squares separate (LSS) approach (Mumford et al., 2012, 2014; Abdulrahman and Henson, 2016) where a separate first-level GLM was specified for each trial of interest within an *a priori* ROI. For instance, the first model included the first fixed payment as a separate regressor while all remaining fixed payment events within the same run were modeled together. Other task-related regressors and nuisance variables were included as in the primary first-level GLM. This processing step produced a beta estimate for each fixed payment, yielding a fixed-payment-specific measure of neural activity while reducing collinearity among regressors.

Using the *mediation* toolbox in Matlab, we performed a multi-level moderated mediation analysis with trial-wise neural estimates within our *a priori* VS ROI to account for the nested structure of trials within participants (Shrout and Bolger, 2002; Woo et al., 2015). Consistent with contemporary approaches (Hayes, 2017), inference focused on the indirect effect (path a X path b), with statistical significance assessed using nonparametric bootstrap resampling with 5000 samples. We constructed a moderated mediation model with participant-specific stimulus (X: trial-wise fixed payment), neural mediator (M: trial-wise neural estimates within VS), outcome behavior (Y: trial-wise effort exertion), and a second-level participant-specific moderator (L2M: participant-specific wage-effort mapping). X, Y, and M were z-scored within each participant. If the first-level indirect effect exists without the direct effect (path c), this suggests a full mediation effect of the neural mediator. Whereas, if it exists with the direct effect, the neural mediator serves as a first-level moderator, indicating a partial mediation from participant-specific stimulus to outcome behavior. The second-level moderator was estimated as the slope of a participant-specific regression of exerted effort on the given payment during Recall; this second-level moderator essentially captures the fidelity with which an individual can exert effort at a target wage-earning amount. This model allowed the indirect effect to vary as a function of these individual differences, in addition to a mediation effect between X and Y.

## Results

After practice, participants had a good understanding of the effort-reward associations and were able to successfully exert to goal earning levels in the Training Phase (Fig 1D; t(118) = 61.44, p < 0.001). The training at these levels transferred to the Recall Phase, such that participants were able to achieve the goal-earning levels without online visual feedback of their exertion trajectory (Fig 1E; t(118) = 22.73, p < 0.001).

### Behavioral Representations of Reference-Dependent Effort Allocation

We hypothesized that participants could predominantly adopt two approaches when deciding how much effort to allocate, given the possibility of receiving either earnings proportional to their exertion or a fixed payment. If a participant felt that the effort was not costly or aversive, they could exert their maximum effort to maximize potential wage earnings and the expected outcome value of a trial, while disregarding the level of the fixed payment (Fig 2A, left panel). In another case, if a participant finds effort costly and aversive and wants to avoid exerting effort that may not lead to a wage rate payment, they could modulate their exertion levels to account for the fixed payment in hopes of reducing the possibility of wasted effort (Fig 2A, right panel). In this way, the amount of effort a participant exerts and their willingness to overshoot or undershoot the fixed payment value with their piece-rate earnings reflect their cost of effort and indicate how the fixed payments may serve as a reference point that motivates their effort provision.

**Fig 2.**
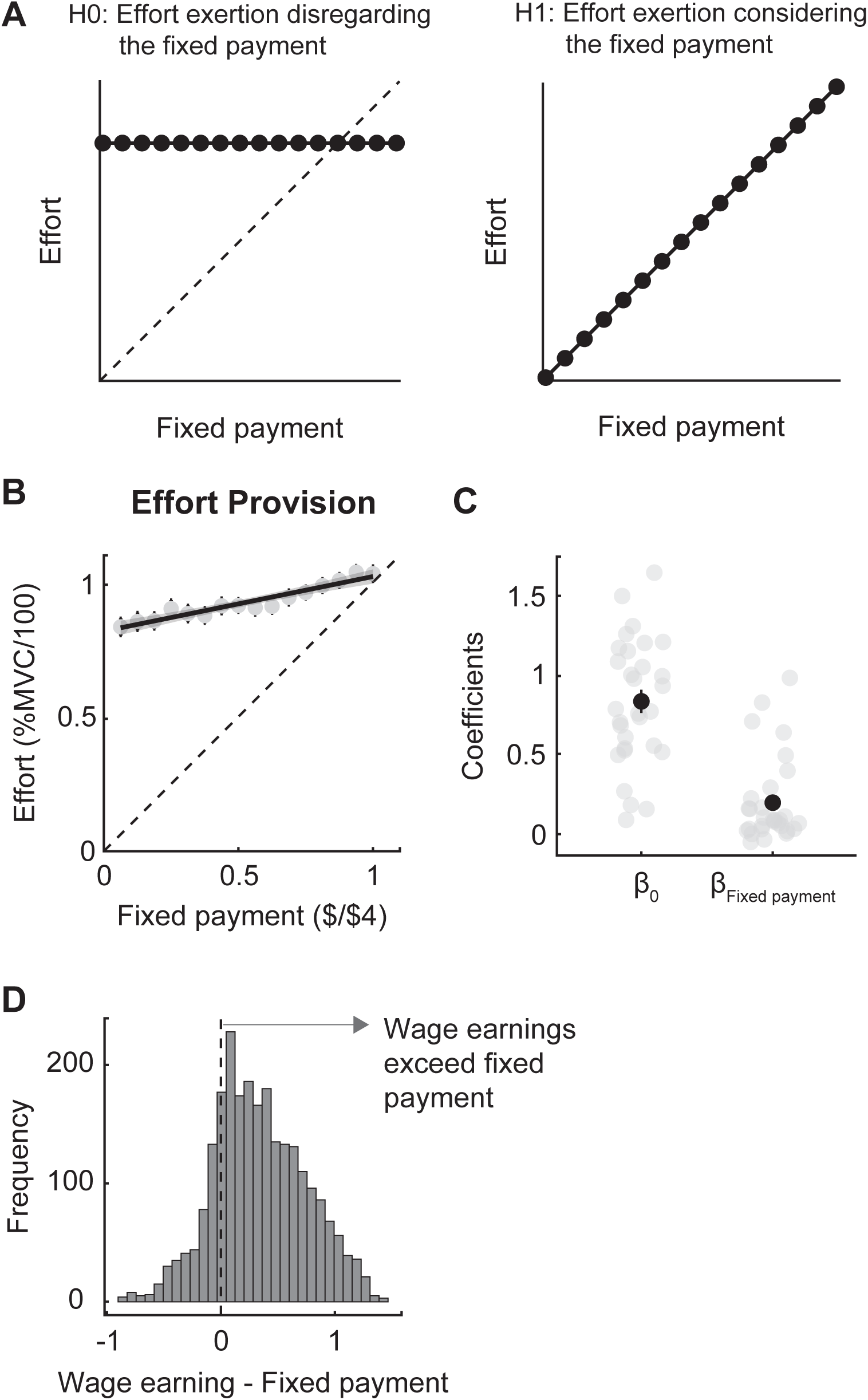
Behavioral Results. Participants could take two approaches in deciding how much effort to allocate when presented with the possibility of receiving either earnings proportional to their exertion or a fixed payment. (**A left**) In one case, if an individual felt that the effort was not costly or aversive, they could exert their maximum effort to maximize potential wage earnings and the expected outcome value of a trial and disregard the level of the fixed payment. (**A right**) In another case, if participants found effort costly and aversive and wanted to avoid exerting effort that might not lead to a wage payment, they could modulate their exertion levels to account for the fixed payment amount, in hopes of reducing their exertion to avoid wasted effort. (**B**) Effort exertion as a function of the fixed payment during the Effort Provision Phase. Both effort and fixed payment were normalized so that 1 effort unit corresponded to 100 effort units, and 1 unit of fixed payment corresponded to $4. The gray points and error bars represent the mean and standard error of effort exertion across participants for each fixed payment. The black line and shaded area represent the regression fit and confidence intervals across all participants. The dotted line represents the case where participants’ wage earnings match the fixed payment. (**C**) Intercept and slope coefficients from the individual regression models. Gray points represent individual participants; the black points and whiskers are the mean and standard error across participants. (**D**) A histogram of how effort-based wage earning deviated from fixed payment. A value greater than 0 indicates wage earnings exceeding the effort required for the fixed payment.

Consistent with a strategy in which the fixed payment modulated exertion levels, we found that participants exerted more effort and increased their potential wage earnings as the fixed payment increased (t(1426) = 13.70, p < 0.001, Fig 2B). There was considerable inter-individual variability in effort provision across fixed payment values (Fig 2C). The majority of participants exhibited a positive intercept and slope in the relationship between the fixed payment and effort exertion (Fig 2C). Moreover, in most trials, the effort provided exceeded the effort required for the trial’s fixed payment (Fig 2D). These results suggest that the cost of exertion, and associated wage earnings, did not exceed the value of the fixed payment. Consistent with previous behavioral studies of reference-dependent effort provision (Fehr and Goette, 2007; Abeler et al., 2011; Crawford and Meng, 2011; Gill and Prowse, 2012; Allen et al., 2014; Bartling et al., 2015; Gutierrez et al., 2021), our results suggest that the fixed payment influences participants’ exogenous expectations of prospective earnings and shapes motivation to exert effort.

### Neural substrates encoding reward expectations from effortful exertion

At the time of incentive presentation, we found that VS activity increased with the value of the fixed payment (Fig 3A, B), consistent with our hypothesis and previous literature showing that the VS plays a crucial role in signaling increasing prospective reward value (Cardinal et al., 2002; Kable and Glimcher, 2007; Levy and Glimcher, 2012; Bartra et al., 2013).

**Fig. 3.**
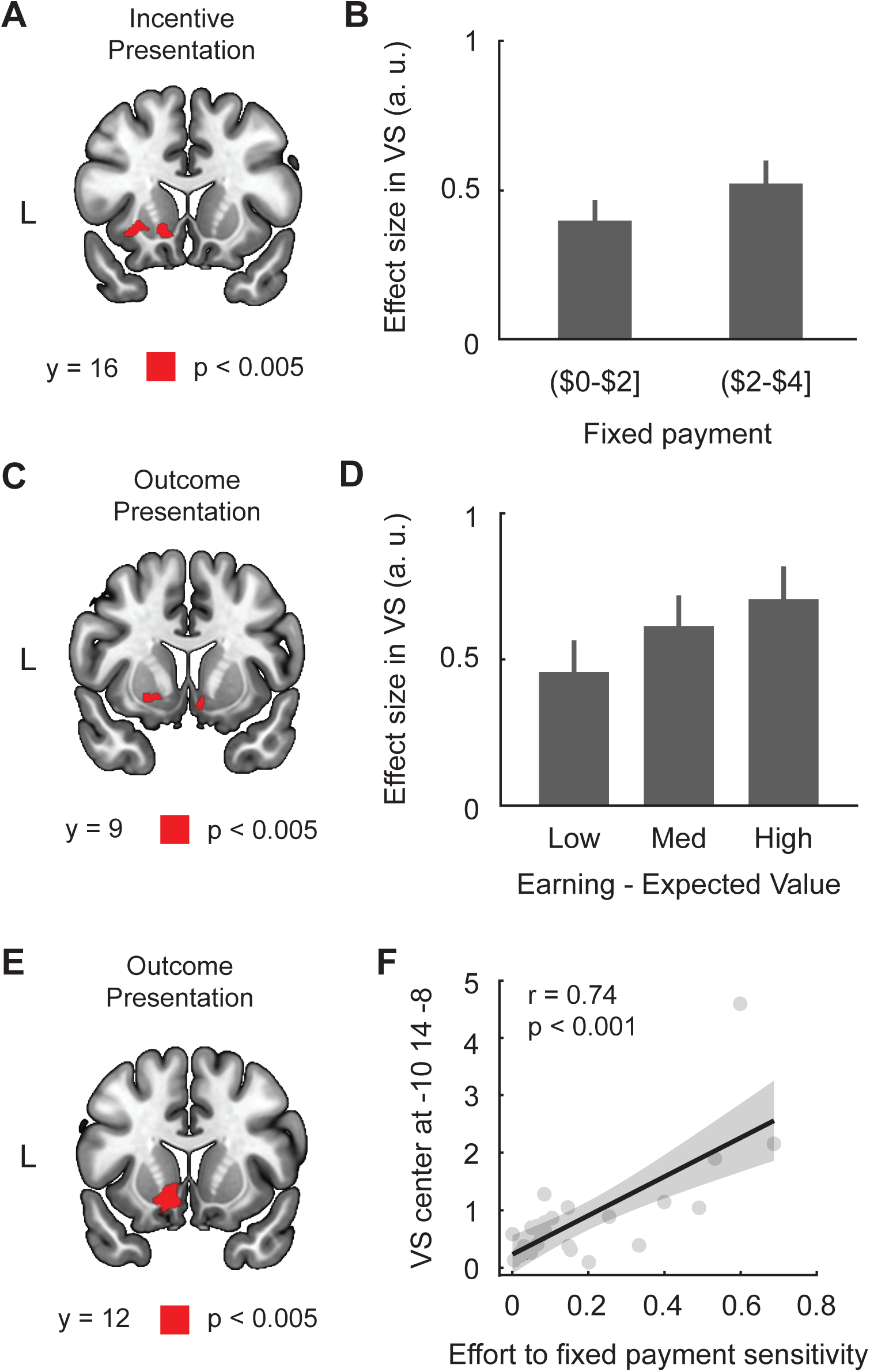
fMRI Results. (**A**) Whole brain results during the Effort Provision Phase at the time of Incentive Presentation. Activity in the left VS (peak = [-16, 10, -6]; small volume corrected p < 0.05 in a priori ROI) was positively modulated by the fixed payment. (**B**) Effects in the right VS (5 mm sphere centered at [-13, 17, -8]) for low ($0.25-$2) and high ($2.25-$4) fixed payments. This was illustrative only and was not used for statistical inference. (**C**) Whole brain results during Outcome Presentation. Activity in the VS (left peak = [-10, 12, -4], right peak = [16, 6, -8], small volume corrected p < 0.05 in a priori ROI) reflected the difference between the actual outcome and the expected value for the fixed payment and wage payment, given an individual’s exertion. (**D**) Effects in the bilateral VS (5mm sphere centered at [-6, 7, -13] for left and [21, 7, -10] for right) for low, medium, and high deviations of the outcome from the reward expectations. This was illustrative only and was not used for statistical inference. (**E**) Whole-brain covariate results during the Outcome Presentation. There was significant activity in the left VS associated with the degree of an individual’s reference-dependent effort exertion (peak = [-10, 14, -8]). (**G**) Neural activity in the nucleus accumbens was significantly correlated with individual differences in reference-dependent tendency during effortful activity, as estimated from the individual regression model (r=0.74, p<0.001).

At the time of outcome presentation, we found that VS activity encoded the difference in value between the actual outcome (fixed payment or wage earnings) and the overall expected value of a trial, given potential earnings from exertion and the fixed payment (0.5*wage earnings + 0.5*fixed payment) (Fig 3C, D). Notably, this signal could be positive if the actual outcome exceeded the expected value for a given trial, and negative if it was below the expected value (Supplementary Figure 2). This modulation of activity with deviation between the actual outcome and reward expectations is consistent with previous findings that the VS processes reward prediction error (Pagnoni et al., 2002; O’Doherty et al., 2003; Knutson and Cooper, 2005; Pessiglione et al., 2006; Martino et al., 2009; Rutledge et al., 2010; Daw et al., 2011). These results suggest that VS encodes expectation-based preferences arising from effortful exertion and illustrate that the ventral striatum encodes a prediction-error-like signal at the time of outcome that reflects reward expectations resulting from effortful exertion.

We reasoned that if VS encodes reference-dependent effort-provision signals at the time of the outcome, this should be reflected in a relationship between participants’ behavioral sensitivity to reference-dependent exertion and the effort they exert. To test this idea, we used participants’ slope relating the fixed payment to effort provision as a metric of behavioral sensitivity to the fixed payment, which served as a reference point for effort provision. Individuals with a higher slope were more sensitive to the need to modulate their exertion in response to the potential fixed payment – a proxy for encoding a reference point for effort provision. To test this prediction, we entered this behavioral metric as a between-participant covariate in the whole-brain analysis of expectation signals at the time of outcome and found that this parameter significantly modulated the response to the difference between the actual outcome and the expected value for a trial (Fig 3E, F). Participants showed a neurometric-psychometric match between their behavioral and neural responses to the encoding of reference-dependent effort provision in response to the offered fixed payment.

### VS activity moderates the relationship between the fixed payment and exerted effort

We next examined whether individual differences in wage-effort mapping moderated the relationship between VS activity and effort exertion. We hypothesized that participants with a more precise understanding of how exerted effort translates into earnings would show a stronger coupling between VS activity and effort provision. Consistent with this prediction, wage-effort mapping significantly moderated the relationship between VS activity and effort exertion (z = 2.92, p = 0.0035; Fig. 4A, B). Participants who more accurately mapped expected earnings onto exerted effort exhibited a stronger association between VS activity and subsequent effort provision based on fixed payment (Fig. 4C). Together, these findings suggest that VS signals contribute to reference-dependent effort behavior most strongly when reward expectations can be reliably translated into exertion.

**Fig. 4.**
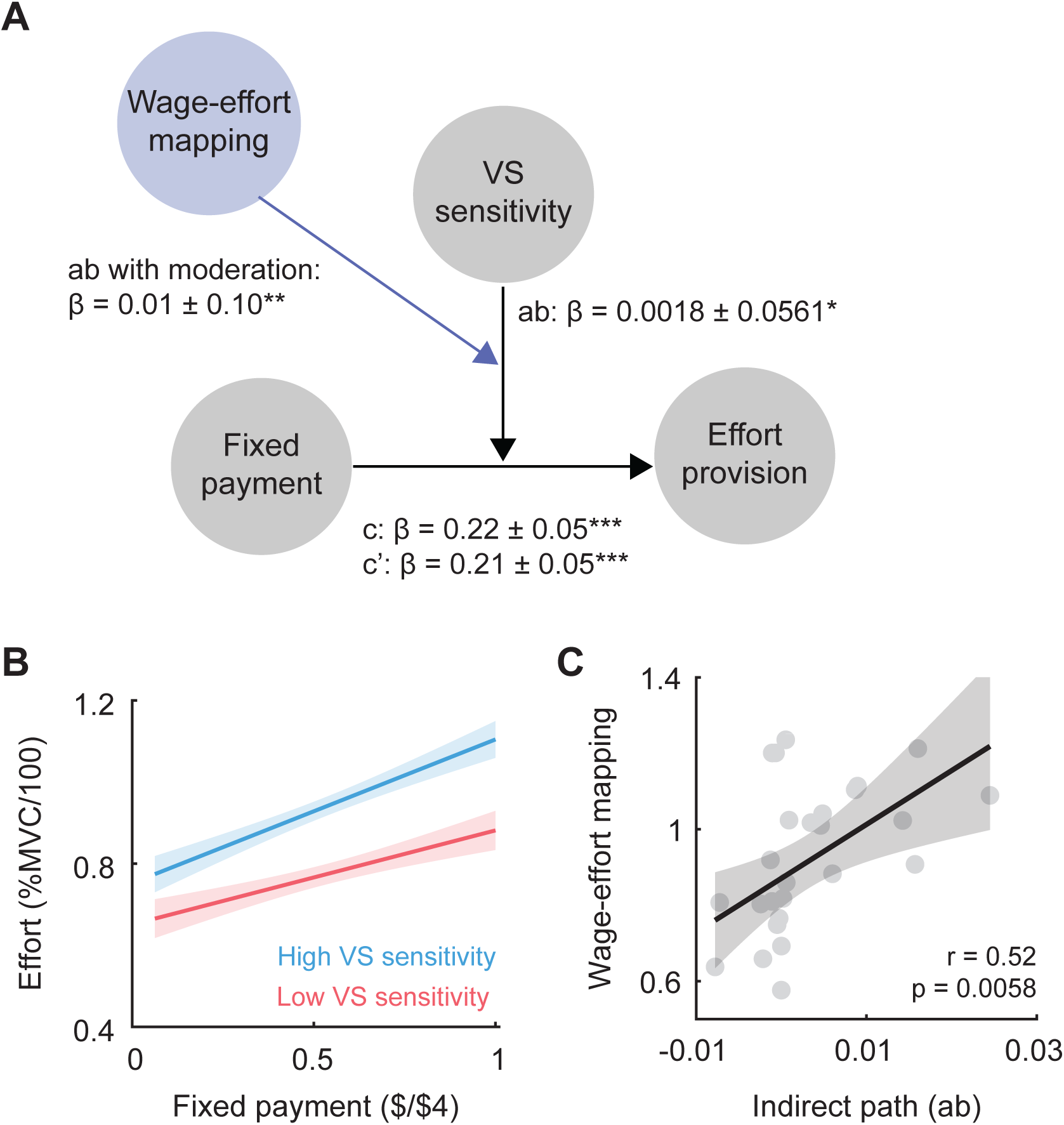
Moderated mediation analysis results. **(A)** Mediation results with added moderation. We constructed a mediation model with trial-wise fixed payment as a predictor, X, trial-wise VS activity as a mediator, M, and trial-wise effort provision as an outcome, Y. There was a significant indirect effect (ab: β = 0.0006, SE = 0.0004, p = 0.04). Statistical results on path a, b, c, and c’ are provided. We further tested the moderation effect of individual differences in the degree of wage-effort mapping. This participant-specific mapping significantly modulated the indirect effect of the mediation model (β = 0.007, SE = 0.0023, p = 0.003). **(B)** A visualization of the first level moderation effect of VS sensitivity on the relationship between fixed payment and effort provision. Participants were split in half based on the indirect path coefficient (High in blue and Low in red). Participants who had a higher moderation effect of the neural estimates showed more sensitivity in fixed payment on effort provision. **(C)** The wage- effort mapping was positively correlated with the indirect path (ab, r = 0.52, p = 0.0058).

## Discussion

Effortful behavior is often conceptualized as a trade-off between reward and cost (De Borger and Fosgerau, 2008; Chong et al., 2017; Müller et al., 2021; Burrell et al., 2023), yet such formulations leave open a fundamental question: relative to what reference point is effort evaluated? Economic theories of reference dependence propose that labor supply and effort respond not to absolute incentives but to deviations from expected or target earnings (Fehr and Goette, 2007; Farber, 2008; Abeler et al., 2011; Crawford and Meng, 2011). However, the neural mechanisms by which reward expectations influence effort remain poorly understood. Here, we provide behavioral and neural evidence that reward expectations serve as a reference point for effort valuation, shaping both effort provision and neural activity during effortful behavior.

By experimentally manipulating reward expectations independently of outcomes, we demonstrate that effort provision is systematically biased by an externally imposed reference point. Participants adjusted their exertion as a function of expected reward, indicating that effort is evaluated relative to anticipated outcomes rather than absolute incentives alone. These findings extend economic theories of reference-dependent labor supply by providing evidence that reward expectations directly influence effort exertion. More broadly, they suggest that reference-dependent preferences are not limited to economic or market decisions but reflect a general principle governing motivated behavior.

At the neural level, activity in the ventral striatum (VS) tracked both reward expectations at the time of choice and deviations between expected and realized outcomes. These signals parallel reference-dependent value computations observed in economic decision-making (Pagnoni et al., 2002; Knutson and Cooper, 2005; Martino et al., 2009), providing a neural substrate through which expectations influence effort.

Importantly, VS activity also explained inter-individual variability in the degree of reference dependence, linking neural valuation processes to behavioral differences. These findings suggest that the VS represents outcomes relative to expected rewards, consistent with its broader role in subjective value computation (Chib et al., 2009; Bartra et al., 2013; Sescousse et al., 2015).

The present findings also extend neurobiological accounts of effort-based decision-making. Much of the existing literature has focused on how effort costs, reward magnitude, and physiological state shape willingness to exert effort (Salamone et al., 2016, 2018; Chong et al., 2017; Müller et al., 2021; Salamone and Correa, 2023). Our previous work has similarly shown that effort valuation is dynamically influenced by internal state signals, including cognitive fatigue, neuromuscular fatigue, and subjective effort perception (Casamento-Moran et al., 2025; Dryzer et al., 2025; Steward et al., 2025, 2026). Together, these studies suggest that the subjective value of exertion is not fixed but is continuously recalibrated according to an individual’s current physiological and cognitive state. The present findings extend this framework by demonstrating that external contextual information—specifically reward expectations—also contributes to effort valuation. Rather than evaluating rewards in absolute terms, individuals assess effort relative to expected outcomes, indicating that motivational value is shaped jointly by internal states and external reference points. These findings provide a bridge between economic theories of reference dependence and neurobiological models of motivated behavior, and suggest that effort allocation emerges from integrating both internal and external sources of valuation.

The relationship between VS activity and effort exertion was further influenced by the efficacy with which participants could translate reward information into exertion. Individuals with a stronger wage-effort mapping showed a stronger association between VS activity and effort provision, indicating that neural representations of expected value do not uniformly influence behavior across individuals. Instead, the impact of VS signals appears to depend on how effectively reward information is transformed into effortful exertion. Consistent with broader models of decision-making, value representations may require effective coupling to motor systems before they can guide behavior (Mogenson et al., 1980; Rangel et al., 2008; Salamone et al., 2016). Together, these findings suggest that reference-dependent effort provision depends on both neural representations of reward expectations and the ability to translate those representations into exertion.

One limitation in comparing our findings with previous labor-supply studies is the differing nature of effort costs. Participants in the present study incurred transient physical effort costs, whereas many prior studies examined decisions involving irreversible temporal costs (Abeler et al., 2011; Gneezy et al., 2017). This distinction may explain why participants, on average, exerted more effort than would be predicted by a strict earnings-equalization strategy. More generally, reference-dependent effort valuation may depend on the perceived recoverability of effort costs. Future studies that directly compare the physical, cognitive, and temporal effort domains may clarify how different cost structures shape expectation-based motivation.

It should be noted that the present study focused on physical effort exertion using a grip-force task, and it remains unclear whether similar reference-dependent mechanisms generalize to cognitive effort. This question is particularly important given accumulating evidence that both cognitive and physical fatigue alter effort valuation and willingness to exert effort (Dryzer et al., 2025; Steward et al., 2025). Determining whether reward expectations interact with fatigue signals to jointly regulate effort allocation represents an important direction for future research.

Together, these findings demonstrate that reward expectations function as reference points for effort exertion and identify ventral striatal signals associated with this process. By linking reference-dependent preferences to neural mechanisms of effort exertion, the present work extends theories of motivated behavior beyond absolute reward-effort trade-offs. More broadly, our findings support the view that effort valuation is a dynamic process shaped not only by anticipated costs and rewards but also by contextual expectations. Understanding how these signals are integrated may provide a unifying framework for explaining adaptive effort allocation across healthy behavior and disorders characterized by impaired motivation.

## Acknowledgments

This work was supported by the Eunice Kennedy Shriver National Institute of Child Health and Human Development and the National Institutes of Health under Award Numbers R01HD097619 and R01MH119086 (VSC).

## Data and materials availability

Data and codes used in the analysis are available in the project’s Open Science Framework website (https://osf.io/t3q72/).

## Author Contributions

Conceptualization: AK, VSC; Methodology: AK, VSC; Investigation: AK, VSC; Visualization: AK, VSC; Funding acquisition: VSC; Project administration: AK, VSC; Supervision: VSC; Writing – original draft: AK; Writing – review & editing: AK, VSC

**Supplementary Figure 1.** Coefficients from the regression model for potential experimental confounds. The bars represent coefficients for each independent variable in the model. Red bars indicate significant effects on effort exertion at trial t. Gray bars indicate no significance.

**Supplementary Figure 2.** A histogram of the parametric modulator (Earning – Expected Value) used in the analysis during the Outcome Presentation.

## References

Abdulrahman H, Henson RN (2016) Effect of trial-to-trial variability on optimal event-related fMRI design: Implications for Beta-series correlation and multi-voxel pattern analysis. NeuroImage 125:756–766.

Abeler J, Falk A, Goette L, Huffman D (2011) Reference Points and Effort Provision. Am Econ Rev 101:470–492.

Allen EJ, Dechow PM, Pope DG, Wu G (2014) Reference-Dependent Preferences: Evidence from Marathon Runners. National Bureau of Economic Research.

Bartling B, Brandes L, Schunk D (2015) Expectations as Reference Points: Field Evidence from Professional Soccer. Manag Sci 61:2646–2661.

Bartra O, McGuire JT, Kable JW (2013) The valuation system: A coordinate-based meta-analysis of BOLD fMRI experiments examining neural correlates of subjective value. NeuroImage 76:412–427.

Burrell M, Pastor-Bernier A, Schultz W (2023) Worth the Work? Monkeys Discount Rewards by a Subjective Adapting Effort Cost. J Neurosci 43:6796–6806.

Camerer C, Babcock L, Loewenstein G, Thaler R (1997) Labor Supply of New York City Cabdrivers: One Day at a Time*. Q J Econ 112:407–441.

Cardinal RN, Parkinson JA, Hall J, Everitt BJ (2002) Emotion and motivation: the role of the amygdala, ventral striatum, and prefrontal cortex. Neurosci Biobehav Rev 26:321–352.

Casamento-Moran A, Kim A, Lee JL, Chib VS (2025) Neuromuscular Signals Shape Fatigue and Effort-Based Decision-Making in Humans. bioRxiv:2025.12.06.692763.

Chib VS, De Martino B, Shimojo S, O’Doherty JP (2012) Neural Mechanisms Underlying Paradoxical Performance for Monetary Incentives Are Driven by Loss Aversion. Neuron 74:582–594.

Chib VS, Rangel A, Shimojo S, O’Doherty JP (2009) Evidence for a Common Representation of Decision Values for Dissimilar Goods in Human Ventromedial Prefrontal Cortex. J Neurosci 29:12315–12320.

Chib VS, Shimojo S, O’Doherty JP (2014) The Effects of Incentive Framing on Performance Decrements for Large Monetary Outcomes: Behavioral and Neural Mechanisms. J Neurosci 34:14833–14844.

Chong TT-J, Apps M, Giehl K, Sillence A, Grima LL, Husain M (2017) Neurocomputational mechanisms underlying subjective valuation of effort costs. PLOS Biol 15:e1002598.

Crawford VP, Meng J (2011) New York City Cab Drivers’ Labor Supply Revisited: Reference-Dependent Preferences with Rational-Expectations Targets for Hours and Income. Am Econ Rev 101:1912–1932.

Croxson PL, Walton ME, O’Reilly JX, Behrens TEJ, Rushworth MFS (2009) Effort- based cost-benefit valuation and the human brain. J Neurosci Off J Soc Neurosci 29:4531–4541.

Daw ND, Gershman SJ, Seymour B, Dayan P, Dolan RJ (2011) Model-Based Influences on Humans’ Choices and Striatal Prediction Errors. Neuron 69:1204– 1215.

De Borger B, Fosgerau M (2008) The trade-off between money and travel time: A test of the theory of reference-dependent preferences. J Urban Econ 64:101–115.

Dryzer MH, Nourbakhsh B, Keller J, Chib VS (2025) Increased Exertion Variability is Linked to Disruptions in Effort Assessment in Multiple Sclerosis. bioRxiv:2025.07.16.665234.

Dunne S, Chib VS, Berleant J, O’Doherty JP (2019) Reappraisal of incentives ameliorates choking under pressure and is correlated with changes in the neural representations of incentives. Soc Cogn Affect Neurosci 14:13–22.

Farber HS (2008) Reference-Dependent Preferences and Labor Supply: The Case of New York City Taxi Drivers. Am Econ Rev 98:1069–1082.

Fehr E, Goette L (2007) Do Workers Work More if Wages Are High? Evidence from a Randomized Field Experiment. Am Econ Rev 97:298–317.

Gill D, Prowse V (2012) A Structural Analysis of Disappointment Aversion in a Real Effort Competition. Am Econ Rev 102:469–503.

Gneezy U, Goette L, Sprenger C, Zimmermann F (2017) The Limits of Expectations- Based Reference Dependence. J Eur Econ Assoc 15:861–876.

Gutierrez C, Obloj T, Frank DH (2021) Better to have led and lost than never to have led at all? Lost leadership and effort provision in dynamic tournaments. Strateg Manag J 42:774–801.

Hayes AF (2017) Introduction to mediation, moderation, and conditional process analysis: A regression-based approach. Guilford publications.

Hogan PS, Chen SX, Teh WW, Chib VS (2020) Neural mechanisms underlying the effects of physical fatigue on effort-based choice. Nat Commun 11:4026.

Hogan PS, Galaro JK, Chib VS (2019) Roles of Ventromedial Prefrontal Cortex and Anterior Cingulate in Subjective Valuation of Prospective Effort. Cereb Cortex 29:4277–4290.

Kable JW, Glimcher PW (2007) The neural correlates of subjective value during intertemporal choice. Nat Neurosci 10:1625–1633.

Kahneman D, Tversky A (1979) Prospect Theory: An Analysis of Decision under Risk. Econometrica 47:263–291.

Knutson B, Cooper JC (2005) Functional magnetic resonance imaging of reward prediction. Curr Opin Neurol 18:411–417.

Kőszegi B, Rabin M (2006) A Model of Reference-Dependent Preferences. Q J Econ 121:1133–1165.

Kurniawan IT, Guitart-Masip M, Dayan P, Dolan RJ (2013) Effort and Valuation in the Brain: The Effects of Anticipation and Execution. J Neurosci 33:6160–6169.

Kurniawan IT, Guitart-Masip M, Dolan RJ (2011) Dopamine and Effort-Based Decision Making. Front Neurosci 5:81.

Levy DJ, Glimcher PW (2012) The root of all value: a neural common currency for choice. Curr Opin Neurobiol 22:1027–1038.

Martino BD, Kumaran D, Holt B, Dolan RJ (2009) The Neurobiology of Reference- Dependent Value Computation. J Neurosci 29:3833–3842.

Mogenson GJ, Jones DL, Yim CY (1980) From motivation to action: functional interface between the limbic system and the motor system. Prog Neurobiol 14:69–97.

Müller T, Klein-Flügge MC, Manohar SG, Husain M, Apps MAJ (2021) Neural and computational mechanisms of momentary fatigue and persistence in effort-based choice. Nat Commun 12:4593.

Mumford JA, Davis T, Poldrack RA (2014) The impact of study design on pattern estimation for single-trial multivariate pattern analysis. NeuroImage 103:130–138.

Mumford JA, Turner BO, Ashby FG, Poldrack RA (2012) Deconvolving BOLD activation in event-related designs for multivoxel pattern classification analyses. NeuroImage 59:2636–2643.

O’Doherty JP, Dayan P, Friston K, Critchley H, Dolan RJ (2003) Temporal difference models and reward-related learning in the human brain. Neuron 38:329–337.

O’Donoghue T, Sprenger C (2018) Reference-Dependent Preferences. In: Handbook of Behavioral Economics: Applications and Foundations 1 (Bernheim BD, DellaVigna S, Laibson D, eds), pp 1–77 Handbook of Behavioral Economics - Foundations and Applications 1. North-Holland.

Pagnoni G, Zink CF, Montague PR, Berns GS (2002) Activity in human ventral striatum locked to errors of reward prediction. Nat Neurosci 5:97–98.

Pessiglione M, Schmidt L, Draganski B, Kalisch R, Lau H, Dolan RJ, Frith CD (2007) How the brain translates money into force: a neuroimaging study of subliminal motivation. Science 316:904–906.

Pessiglione M, Seymour B, Flandin G, Dolan RJ, Frith CD (2006) Dopamine-dependent prediction errors underpin reward-seeking behaviour in humans. Nature 442:1042–1045.

Power JD, Mitra A, Laumann TO, Snyder AZ, Schlaggar BL, Petersen SE (2014) Methods to detect, characterize, and remove motion artifact in resting state fMRI. NeuroImage 84:10.1016/j.neuroimage.2013.08.048.

Prévost C, Pessiglione M, Météreau E, Cléry-Melin M-L, Dreher J-C (2010) Separate Valuation Subsystems for Delay and Effort Decision Costs. J Neurosci 30:14080–14090.

Rangel A, Camerer C, Montague PR (2008) A framework for studying the neurobiology of value-based decision making. Nat Rev Neurosci 9:545–556.

Rutledge RB, Dean M, Caplin A, Glimcher PW (2010) Testing the Reward Prediction Error Hypothesis with an Axiomatic Model. J Neurosci 30:13525–13536.

Salamone J, Correa M (2023) The Neurobiology of Activational Aspects of Motivation: Exertion of Effort, Effort-Based Decision Making, and the Role of Dopamine. Annu Rev Psychol 75.

Salamone JD, Correa M, Yang J-H, Rotolo R, Presby R (2018) Dopamine, Effort-Based Choice, and Behavioral Economics: Basic and Translational Research. Front Behav Neurosci 12.

Salamone JD, Pardo M, Yohn SE, López-Cruz L, SanMiguel N, Correa M (2016) Mesolimbic Dopamine and the Regulation of Motivated Behavior. In: Behavioral Neuroscience of Motivation (Simpson EH, Balsam PD, eds), pp 231–257 Current Topics in Behavioral Neurosciences. Cham: Springer International Publishing.

Sescousse G, Li Y, Dreher J-C (2015) A common currency for the computation of motivational values in the human striatum. Soc Cogn Affect Neurosci 10:467– 473.

Shrout PE, Bolger N (2002) Mediation in experimental and nonexperimental studies: New procedures and recommendations. Psychol Methods 7:422–445.

Steward G, Looi V, Chib VS (2025) The Neurobiology of Cognitive Fatigue and Its Influence on Effort-Based Choice. J Neurosci Off J Soc Neurosci 45:e1612242025.

Steward GE, Culbreth AJ, Goes FS, Chib VS (2026) Fatigue Symptoms Influence Effort-Based Decision-Making in Major Depressive Disorder. Biol Psychiatry Glob Open Sci:100780.

Tom SM, Fox CR, Trepel C, Poldrack RA (2007) The Neural Basis of Loss Aversion in Decision-Making Under Risk. Science 315:515–518.

Woo C-W, Roy M, Buhle JT, Wager TD (2015) Distinct Brain Systems Mediate the Effects of Nociceptive Input and Self-Regulation on Pain. PLOS Biol 13:e1002036.

Yarkoni T, Poldrack RA, Nichols TE, Van Essen DC, Wager TD (2011) Large-scale automated synthesis of human functional neuroimaging data. Nat Methods 8:665–670.

