## Supplemental Figures for "Neural substrates underlying the expectation of rewards resulting from effortful exertion"

**Affiliations:**

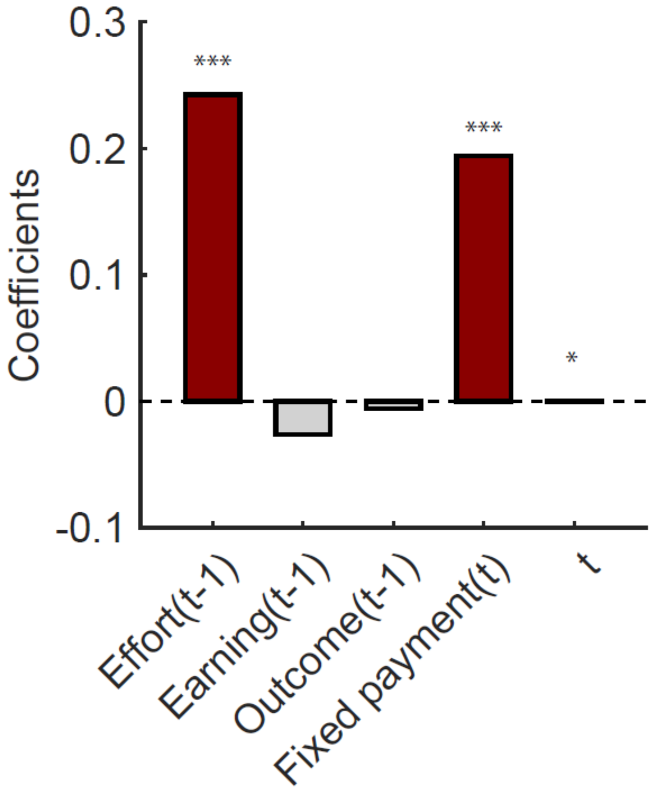


**Fig. S1.**

Coefficients from the regression model for potential experimental confounds. The bars represent coefficients for each independent variable in the model. Red bars indicate significant effects on effort exertion at trial t. Gray bars indicate no significance.

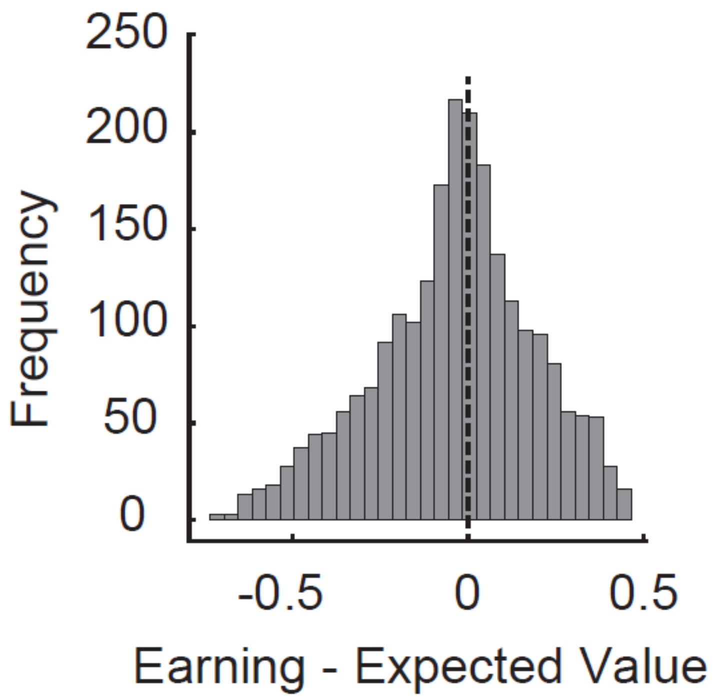


**Fig. S2.**

A histogram of the parametric modulator (Earning – Expected Value) used in the analysis during the Outcome Presentation.
